# Adaptive eQTLs reveal the evolutionary impacts of pleiotropy and tissue-specificity, while contributing to health and disease in human populations

**DOI:** 10.1101/444737

**Authors:** Melanie H. Quiver, Joseph Lachance

## Abstract

Regulatory DNA has the potential to be adaptive, and large numbers of expression quantitative trait loci (eQTLs) have recently been identified in humans. For the first time, a comprehensive study of adaptive eQTLs is possible. Many eQTLs have large allele frequency differences between populations, and these differences can be due to natural selection. Here, we combined population branch statistics with tissue-specific eQTL data to identify positively selected loci in human populations. Adaptive eQTLs tend to affect fewer tissues than non-adaptive eQTLs. Because the tissue breadth of an eQTL can be viewed as a measure of pleiotropy, these results suggest that pleiotropy can inhibit adaptation. The proportion of eQTLs that are adaptive varies by tissue, and we find that eQTLs that regulate expression in testis, thyroid, blood, or sun-exposed skin are enriched for adaptive outliers. By contrast, eQTLs that regulate expression in the cerebrum or female-specific tissues have a relative lack of adaptive outliers. These results reveal tissues that have been the targets of adaptation during the last 100,000 years of human evolution. The strongest adaptive signal in many regions of the human genome is an eQTL, including an eQTL associated with the Duffy blood group and malaria resistance. Scans of selection also reveal that many adaptive eQTLs are closely linked to disease-associated loci. Taken together, our results indicate that adaptive eQTLs have played an important role in human evolution.

## Introduction

Regulatory mutations and changes in gene expression lead to functional differences in anatomy, physiology, and behavior that are evolutionarily important [1-6]. Polymorphic sites that influence gene expression are known as expression quantitative trait loci (eQTLs), and many of these sites are relevant to human health and disease [7-9]. Although identifying specific nucleotides that cause differences in gene expression can be challenging [10], many eQTLs have been identified in model and non-model organisms [11-13]. In recent years, hundreds of thousands of human eQTLs have been cataloged in the GTEx (Genotype-Tissue Expression project) and RegulomeDB databases [14-17]. Many of these eQTLs act in a tissue-specific manner, and by studying adaptive eQTLs it is possible to identify the tissues that have been the primary targets of recent human evolution. Although eQTL effect sizes and directions of effect tend to be conserved among human populations [18], array and sequence data reveal that gene expression patterns vary across populations [19, 20] (but see [21]). Many eQTLs have divergent allele frequencies across populations, and local adaptation may underlie these differences [22, 23]. Hereditary disease risks have evolved in the recent past [24], and many of these changes are likely to be due to positive selection acting on regulatory DNA.

Adaptation is a fundamental concern of evolutionary biology, and recent years have seen a contentious debate about whether adaptation tends to proceed via nonsynonymous changes in coding regions (amino acid changes) or due to changes in gene regulation [25-27]. Of particular relevance is the fact that less than only 1.5% of the human genome is coding [28], and many scans of positive selection have implicated intergenic regions of the human genome [29, 30]. The functional consequences of Neanderthal introgression also tend to involve changes in gene expression, as opposed to protein changes [31]. Regardless of the proportion of the human genome that is functional [32, 33], there are multiple reasons why changes in gene expression may be beneficial. Unlike nonsynonymous changes that affect protein sequences across the body, eQTLs can modify gene expression in a tissue-specific manner. The amount and timing of gene expression can also be optimized for a given environment [34]. Many methods of detecting positively selected alleles exist, including population branch statistics (PBS) [35-37]. These within-species scans of selection use genetic distances between multiple populations to identify outlier loci that have undergone accelerated evolution along one branch of a population-level phylogenetic tree. Scans of selection that examine population differentiation, such as PBS, are well-suited for detecting selection that has occurred on a continental scale during the last 100,000 years [38]. Because PBS scores do not rely on extended haplotype heterozygosity, they are robust to whether adaptive alleles are due to new mutations or standing genetic variation. PBS scores are also able to detect partial sweeps.

Evolutionary theory informs our understanding about which types of eQTLs are expected to be adaptive. In the Fisher-Orr geometric model, adaptation tends to proceed via progressively smaller changes [39-41] (but see [42]). Because of this, we predict that adaptive eQTLs are unlikely to involve large changes in gene expression. Similarly, pleiotropy can inhibit adaptation [43], which leads to the prediction that most adaptive eQTLs will affect a small number of tissues. Scans of selection in human genomes have revealed that immunity genes tend to be fast evolving [44, 45]. There is also evidence from *Drosophila* that testis-expressed genes evolve quickly [46], and reproductive genes have experienced elevated rates of evolution in many vertebrate lineages [47]. Because of this, eQTLs that affect fast-evolving tissues are expected be enriched for adaptive PBS outliers. Despite these predictions, multiple knowledge gaps exist. The extent to which eQTL effect sizes and tissue breadth constrain human adaptation has yet to be tested empirically. It is also unknown whether tissues that have been targets of recent human adaptation are the same tissues that experienced accelerated evolution over deeper timescales. Importantly, affordable sequencing has ushered in an era of population genomics, and thousands of tissue-specific eQTLs have recently been identified [16]. For the first time, a comprehensive understanding of adaptive eQTLs in human populations is possible.

Here, we combine continental allele frequencies from the 1000 Genomes Project with eQTL data from the GTEx project and RegulomeDB to identify adaptive eQTLs in human populations. We focus on five questions: 1) Which eQTLs exhibit signatures of local adaption? 2) Are pleiotropic eQTLs less likely to be positively selected? 3) Does the effect size of an eQTL affect whether it is adaptive? 4) Which tissues tend to be targets of local adaptation? 5) To what extent do adaptive eQTLs overlap with GWAS results?

## Results

### Scans of selection identify adaptive eQTLs

Many presently known eQTLs have large allele frequency differences between populations, and some of these eQTLs are targets of local adaptation. V7 GTEx eQTLs have a mean F_ST_ of 0.113 between African and European populations, 0.131 between African and East Asian populations, and 0.090 between European and East Asian populations. We calculated PBS scores for individuals from Africa, Europe, and East Asia at every biallelic SNP in the 1000 Genomes Project (Figure 1). We then identified the top 1% of all V7 GTEx eQTLs with respect to PBS for each population. Because many eQTLs are closely linked, we pruned this set of high PBS eQTLs to eliminate the effects of linkage disequilibrium. After LD-pruning, we found 614 eQTLs that are adaptive outliers for Africa, 561 eQTLs that are adaptive outliers for Europe, and 524 eQTLs that are adaptive outliers for East Asia. Note that PBS outliers for the African branch also include SNPs that experienced large allele frequency changes after the out-of-Africa migration, but prior to the split of European and Asian lineages. In Figure 1, LD-pruned adaptive eQTLs are represented by filled red circles, other eQTLs are represented by open black circles, and 1000 Genomes Project SNPs that are not eQTLs are represented by open gray circles.

**Figure 1.**
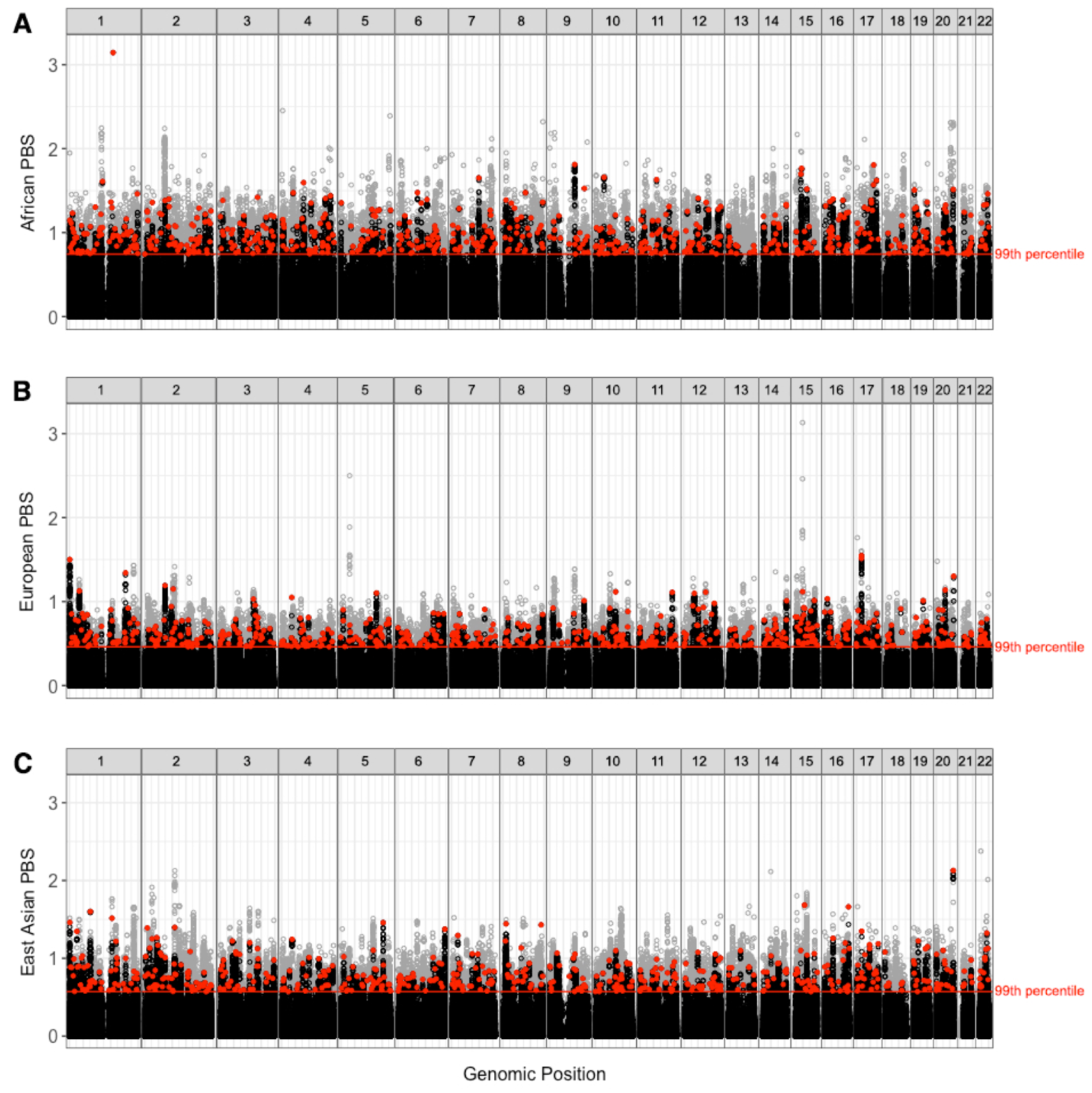
Genome-wide scans of selection identify adaptive eQTLs in human populations. Manhattan plot of Population branch statistics (PBS) vs. genomic position for each population: (A) Africa, (B) Europe, and (C) East Asia. All variants in the 1000 Genomes Project, including V7 GTEx eQTLs, are shown here. LD-pruned adaptive eQTLs are represented by filled red circles, other eQTLs are represented by open black circles, and 1000 Genomes Project SNPs that are not eQTLs are represented by open gray circles.

Scans of selection reveal that the strongest PBS signal in many adaptive regions of the genome is an eQTL (visualized as red-tipped peaks in the Manhattan plots of Figure 1). Overall, we find that GTEx eQTLs are 2.53 times as likely than random SNPs from the 1000 Genomes Project to have signatures of positive selection, i.e. PBS scores above the red lines in Figure 1 (p-value < 0.0001, chi-square test of independence). This underscores the importance of regulatory DNA to recent human evolution. Many adaptive eQTLs are not in linkage disequilibrium with other sites (visualized as isolated red points with high PBS scores in Figure 1), while other putatively adaptive eQTLs appear to cluster in genomes (visualized as red and black points forming a peak in Figure 1). When positive selection acts on standing genetic variation there is enough time for recombination to break down LD between an adaptive allele and linked sites [48]. This can result in an adaptive peak that has only a single high-PBS site. By contrast, when positive selection acts on new mutations a single haplotype can increase in frequency. This results in adaptive peaks that have multiple high PBS sites.

### Pleiotropy and the tissue breadth of eQTLs

Some eQTLs modify gene expression in a small number of tissues, while other eQTLs modify gene expression in many tissues. This can influence whether an eQTL is adaptive. eQTLs analyzed in this present study affect between 1 and 48 tissues, and tissue breadth (the number and types of tissues that each eQTL affects) can be viewed as a measure of pleiotropy. Here, we compare the number of tissues affected by adaptive and non-adaptive eQTLs. Overall, adaptive eQTLs affect fewer tissues than non-adaptive eQTLs. This pattern occurs whether a SNP is adaptive in Africa, Europe, or East Asia (Figure 2). GTEx eQTLs affect a mean number of 5.97 tissues. By contrast, adaptive outliers affect a mean number of 4.08, 4.11 and 3.84 tissues (African, European, and East Asian outliers, respectively). These differences between adaptive outliers and other eQTLs are statistically significant (p-value < 2.2 10-16 for all comparisons, Wilcoxon rank sum tests). Looking beyond the mean number of tissues, we find that adaptive outliers have cumulative distributions that are left-shifted, i.e. they tend to affect a smaller number of tissues (Figure 2). Many non-adaptive eQTLs modify expression in more than 10 tissues, while few adaptive eQTLs modify expression in more than 10 tissues. Furthermore, 55.3% of all LD-pruned adaptive outliers affect a single tissue. Similar patterns arise if a more stringent PBS score cutoff is used (Figure S1): roughly half (50.8%) of eQTLs that have PBS scores above the 99.9th percentile affect a single tissue. Taken together, our results indicate that pleiotropy appears to inhibit adaptation.

**Figure 2.**
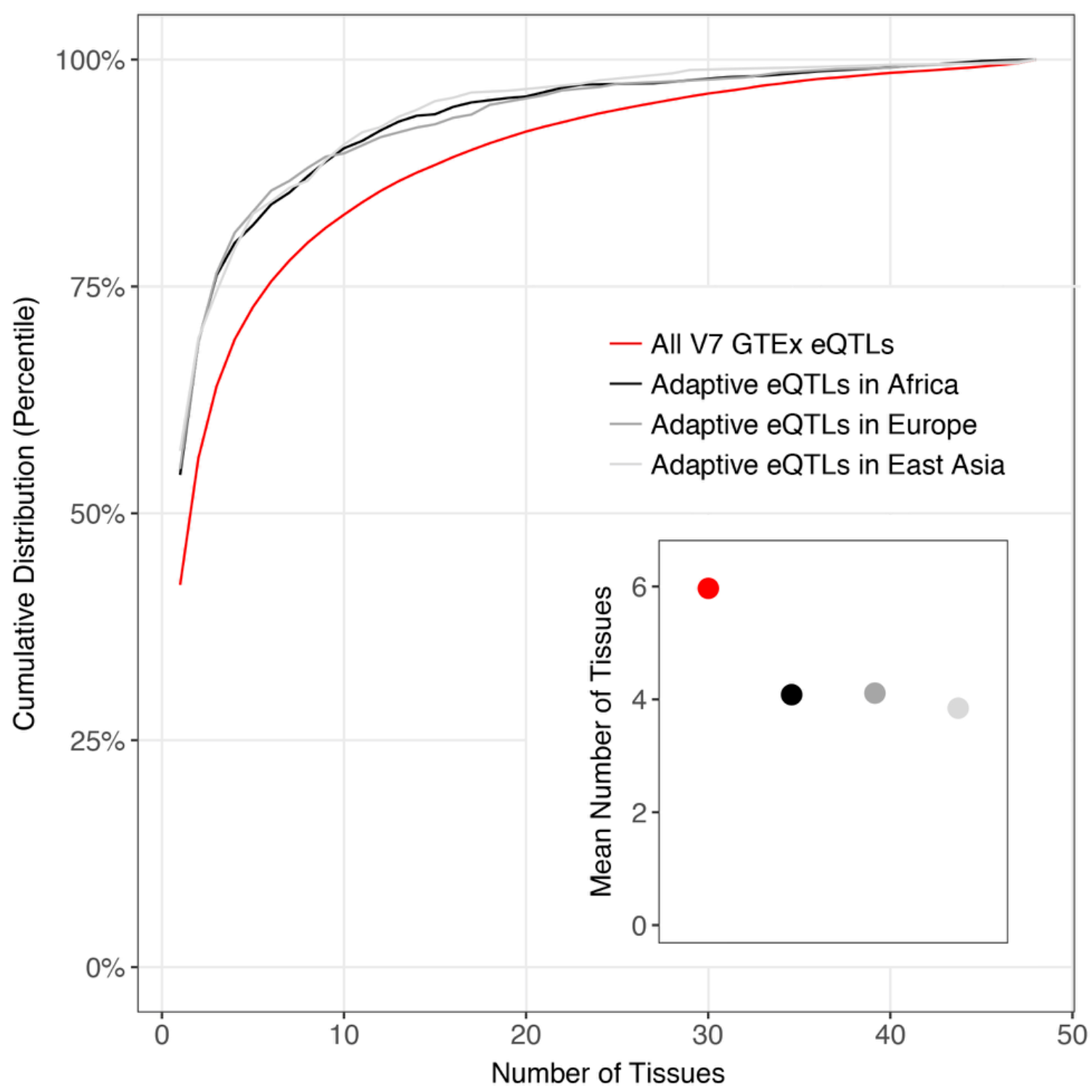
Highly pleiotropic eQTLs are less likely to be adaptive. Cumulative distributions and mean number of tissues are shown for adaptive and non-adaptive eQTLs. Here, adaptive outliers are LD-pruned eQTLs that have PBS scores in the top 1% of all GTEx eQTLs. In general, adaptive eQTLs modify gene expression in fewer tissues than non-adaptive eQTLs. For each population, differences in the number of tissues affected by adaptive outliers and the overall set of GTEx eQTLs are statistically significant (p-value < 2.2 10^-16^ for all comparisons, Wilcoxon rank sum tests).

### Effect sizes and local adaptation

The amount that an allele affects gene expression can potentially influence whether it is adaptive. For each tissue, we compared PBS statistics to the absolute value of normalized effect sizes under a fixed effect model. This allowed us to test whether adaptive eQTLs cause larger changes in expression than non-adaptive eQTLs. This set of analyses focused on LD-pruned eQTLs that only affect a single tissue. In general, adaptive eQTLs do not yield large changes in gene expression. For example, there is a weak negative correlation between PBS scores and effect sizes for eQTLs that modify expression in sun-exposed skin (Figures 3A, 3B and 3C). These patterns occur regardless of whether PBS statistics are calculated for African, European, or East Asian individuals. Examining each of the tissues analyzed in our study, correlations between PBS and |β_FE_| values tend to be close to -0.2 (Figure 3D). Exceptions to this pattern arise for tissues that have a small number of LD-pruned single-tissue eQTLs (gray shading). For each combination of tissue and population we tested whether adaptive PBS outliers have different effect sizes than other eQTLs (Figure 3E). After correcting for multiple tests, we find that 143 out of 144 tissue-population combinations do not yield statistically significant differences in effect sizes between adaptive eQTLs and other eQTLs (Wilcoxon rank sum tests using Bonferroni corrections). Note that eQTLs with very small effect sizes (i.e. |β_FE_| close to zero) are unlikely to be identified in studies like GTEx. Similarly, eQTLs with negligible effect sizes are unlikely to be direct targets of selection. The Fisher-Orr model of adaptation posits that large phenotypic changes are less likely to be adaptive than small changes. Because we observe few adaptive eQTLs with large effect sizes and find weak negative correlations in Figure 3D, our results are consistent with the Fisher-Orr model.

**Figure 3.**
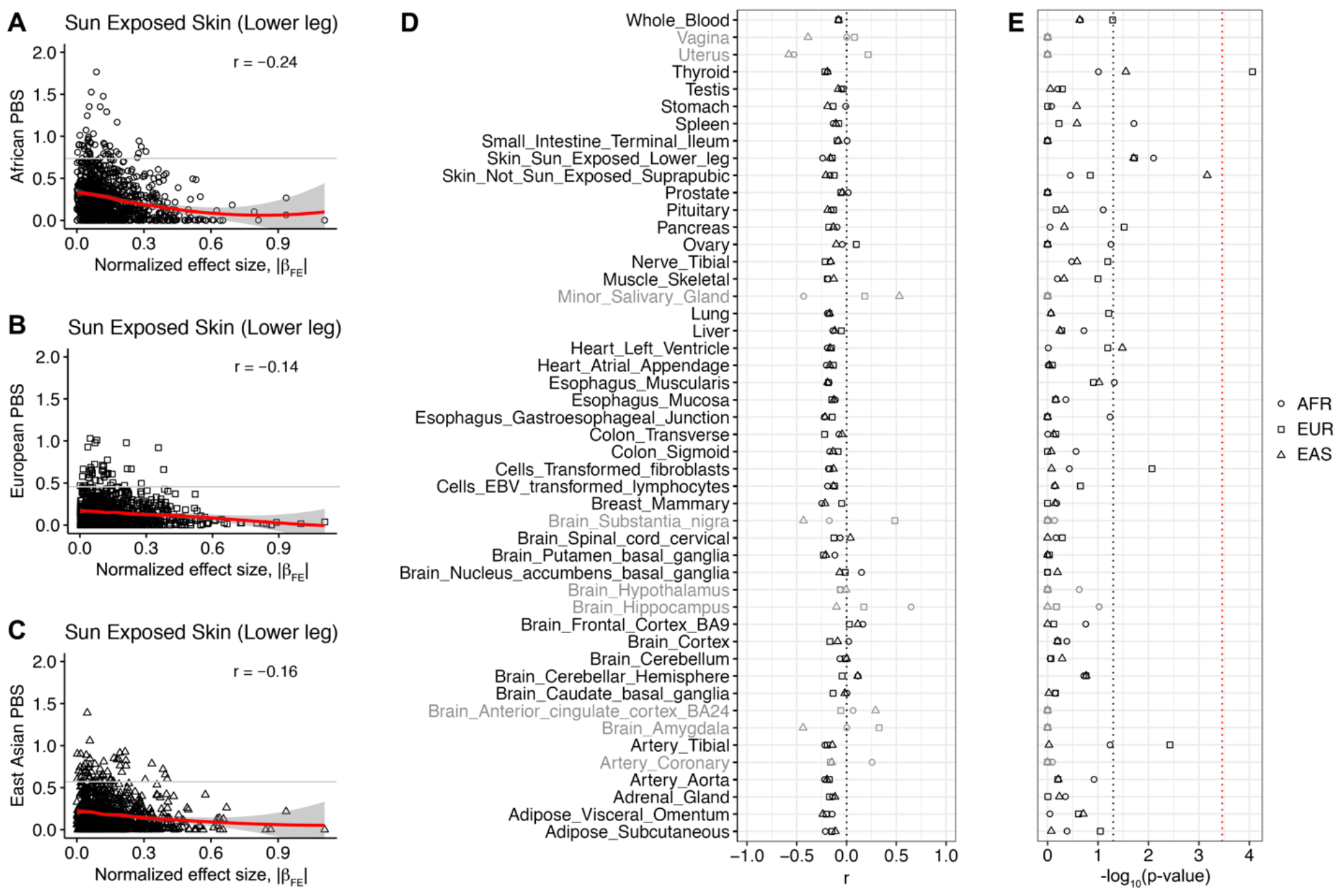
Most adaptive eQTLs do not have a large effect on gene expression. Plots of PBS score vs. |β_FE_| are shown. Each data point represents an LD-pruned SNP that only modifies gene expression in a single tissue, i.e. eQTLs that affect multiple tissues were omitted from this analysis. eQTLs that modify expression in skin exposed to the sun are shown in panels (A), (B), and (C). Curve fitting of scatter plots uses local regression (indicated by red lines). 99^th^ percentiles of PBS scores are indicated by horizontal gray lines in each scatter plot. Population-specific correlations for each tissue are shown in panel (D). Results of Wilcoxon rank sum tests comparing the effect sizes of adaptive and non-adaptive eQTLs are shown in panel (E), with uncorrected (p-value < 0.05) and Bonferroni corrected p-value cutoffs indicated by dotted black and red lines. Gray shading indicates tissues that have fewer than 200 LD-pruned single-tissue eQTLs.

### Tissue-specificity of adaptive eQTLs

The proportion of eQTLs that are adaptive varies by tissue. Here, we used enrichment ratio scores to compare the observed and expected counts of adaptive PBS outliers for each tissue. Positive enrichment ratio scores indicate tissues that have an excess of adaptive eQTLs and negative enrichment ratio scores indicate tissues that have a relative lack of adaptive eQTLs. Because enrichment ratio scores use a natural log scale, a difference of one unit translates to a 2.718-fold difference in the relative proportion of eQTLs that are adaptive outliers.

Focusing on individual tissues, testis eQTLs are the most likely to have high PBS scores, followed by eQTLs that modify gene expression in the thyroid, whole blood, and sun-exposed skin (Figure 4A). Our results suggest that recent human adaptation may have been driven by sexual selection, metabolism, pathogens, and local environmental conditions. We note that adipose, pancreas, and liver are moderately enriched for adaptive eQTLs - indicating that diet has also had an evolutionary impact. Focusing on sex-specific tissues, we find that testis eQTLs are enriched for adaptive PBS outliers. By contrast, we find that eQTLs that affect expression in the prostate, ovary, uterus or vagina have a relative lack of adaptive outliers. We also find that eQTLs that affect expression in the cerebellum are more likely to be adaptive than eQTLs that affect expression in the cerebrum. Pleiotropy contributes to tissue-specific differences in enrichment ratios. Tissues with high enrichment ratios tend to have eQTLs that affect a small number of additional tissues, and tissues with low enrichment ratios tend to have eQTLs that affect many other tissues (Figure 4B). For example, testis eQTLs that are adaptive tend to affect only a small number of additional tissues, if at all (Figures S3, S4, and S5).

**Figure 4.**
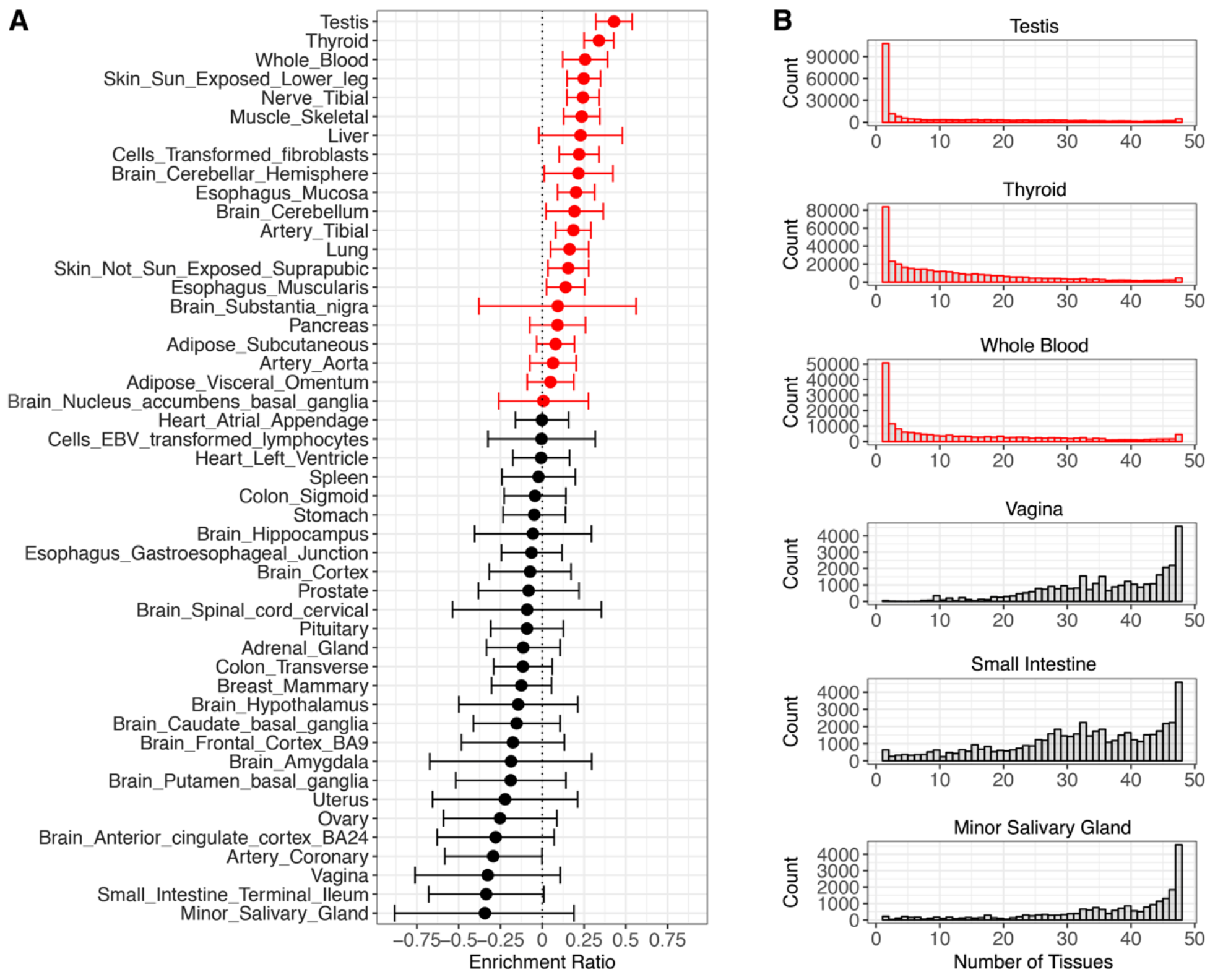
Tissue enrichment of adaptive eQTLs. Positive enrichment ratio scores indicate a relative excess of adaptive eQTLs for a particular tissue (red), and negative enrichment ratio scores indicate a relative lack of adaptive eQTLs for a particular tissue (black). (A) Tissues are ranked by enrichment ratio score. Here, adaptive outliers are LD-pruned eQTLs that have PBS scores in the top 1% of all GTEx eQTLs. 95% confidence intervals for each enrichment ratio are shown. (B) Tissues with high enrichment ratios tend to have eQTLs that affect less than 10 tissues, and tissues with low enrichment ratios tend to have eQTLs that affect more than 40 tissues.

A large number GTEx eQTLs affect gene expression in thyroid tissue, tibial nerves, and sun-exposed skin (Figure S6). One implication of this is that some tissues may be common targets of adaptation simply because they have more eQTLs than other tissues. We also note that a number of tissues that have large enrichment ratios also have large sample sizes. Despite differences in sample sizes between GTEx V6 and GTEx V7, both of these versions of GTEx yield similar enrichment ratios (Figure S2). To further correct for sample size as a covariate, we linearly regressed tissue-specific enrichment ratios against sample size (Figure S7), yielding a set of adjusted enrichment ratios (Figure S8). Fourteen tissues that have both positive enrichment ratios and positive adjusted enrichment ratios (i.e. they are above both the dotted black line and the solid red line in Figure S7). These tissues include testis, thyroid, liver, whole blood, transformed fibroblasts, and the cerebellum. It is also worth noting that there is no single optimal way to correct for sample size (nonlinear regression yields different adjusted enrichment ratios than linear regression).

### eQTLs with the highest PBS scores

Here, we highlight the strongest signatures of adaptation for each population (Figure 5). Each of these eQTLs is an ancestry informative marker. rs2814778 has the highest PBS score for the African branch. This C/T SNP has a striking geographic pattern: African frequencies of the C allele are >96% and non-African allele frequencies of the C allele are <1%. rs2814778 affects gene expression of the *DARC* gene, also known as *ACKR1*. rs2814778 is in the promoter of the *DARC* gene, and the C allele at this regulatory locus confers a null phenotype [49]. *DARC* encodes the Duffy blood group antigen which is known to be adaptive with respect to *Plasmodium vivax* and malaria [50, 51]. rs2814778 is also predictive of neutrophil counts in African-Americans [52]. Despite a lack of local recombination hotspots, rs2814778 has negligible amounts of linkage disequilibrium with nearby SNPs. This hints that selection acting on the Duffy blood group may have acted on standing genetic variation [50]. rs2814778 only modifies expression in whole blood, and this tissue-specificity and lack of pleiotropy may contribute strong signatures of positive selection at this eQTL.

**Figure 5.**
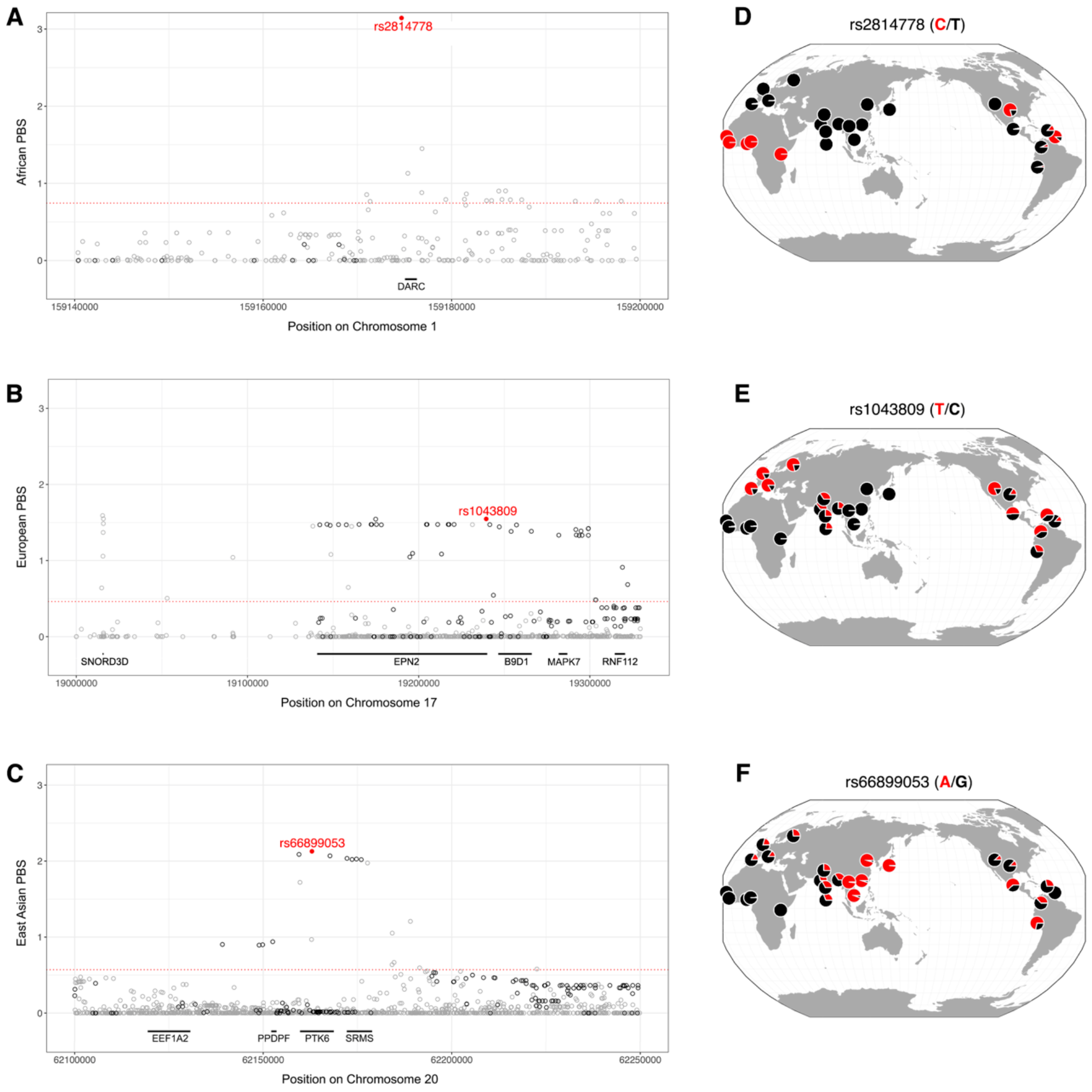
Adaptive eQTLs with strong signatures of positive selection. Genomic regions flanking eQTLs with the highest PBS score for each continental population are shown panels (A), (B), and (C). LD-pruned adaptive eQTLs are represented by filled red circles, other eQTLs are represented by open black circles, and 1000 Genomes Project SNPs are represented by open gray circles. hg19 coordinates are shown. Population-specific 99^th^ percentiles of PBS scores are represented by dashed red lines. Panels (D), (E), and (F) show allele frequencies for 26 populations from the 1000 Genomes Project (modified from the Geography of Genetic Variants Browser [85]).

rs1043809 is the eQTL with the highest PBS score for the European branch. This C/T SNP is near the *EPN2*, *B9D1*, and *RNF112* genes at 17p11. At present, the reason why this genomic region was positively selected in Europe is unknown. Many eQTLs that are closely linked to rs1043809 have similar PBS statistics (visualized as a plateau of points in Figure 5B), which suggests the existence of an adaptive haplotype, rather than a single SNP.

rs66899053 is the eQTL with the highest PBS score for the East Asian branch. This A/G SNP modifies expression of the *EEF1A2, PPDPF*, *PTK6*, and *SRMS* genes, and it is found in an adaptive haplotype at 20q13. Scans of selection have previously implicated this genomic region with respect to *Helicobacter pylori* infection and gastric cancer [53]. Consistent with this cause, rs66899053 affects gene expression in the stomach and many other tissues. Intriguingly, rs66899053 is found in a genomic region that has previously been shown to contain adaptively introgressed Neanderthal alleles in non-African populations [54, 55]. rs66899053 is also 427kb away from HAR1, a genomic region that has undergone accelerated evolution in humans following the split between human and chimpanzee lineages [56].

### Overlap with GWAS results

We also tested the extent to which adaptive outliers overlap with loci that are associated with complex traits and disease susceptibility. An important consideration is that GWAS loci tag genomic regions that are associated with complex traits, i.e. they are sentinel SNPs [57]. Causal SNPs are often 10-100kb distant from sentinel SNPs [58]. A total of 29 adaptive eQTLs are GWAS loci from the NHGRI-EBI Catalog (one shared, six African, twelve European, and ten East Asian PBS outliers). Traits associated with these adaptive eQTLs include white blood cell count, schizophrenia risk, BMI, and eye color. Many eQTLs are closely linked to GWAS loci. Indeed, the median distance of V7 GTEx eQTLs to GWAS loci is 26.4kb. Adaptive PBS outliers have a similar median distance to GWAS loci (26.9kb). By contrast, random SNPs from the 1000 Genomes Project have a median distance to GWAS loci of 30.2kb. These differences are statistically significant: p-value < 2.2 x 10-16 for all eQTLs compared to 1000 Genomes Project SNPs, and p-value = 7.6 x 10-4 for adaptive PBS outliers compared to 1000 Genomes Project SNPs (Wilcoxon rank sum tests). Colocalization of eQTLs and GWAS signals has previously been used to prioritize target genes that are associated with complex traits [59]. Although close proximity to GWAS loci need not imply that regulatory variants are causal, physical linkage can have implications for health inequities [60]. This is because local adaptation results in large allele frequency differences between populations for not only the direct targets of selection, but also linked loci [61]. The combination of positive selection acting eQTLs and genetic hitchhiking may contribute to population-level differences in functionally important traits, including hereditary disease risks.

## Discussion

Our study focused on general patterns of adaptive evolution tissue-specificity, rather than on individual populations. We found that highly pleiotropic eQTLs are less likely to be adaptive than eQTLs that affect a small number of tissues. We also identified tissues that are enriched for adaptive outliers. A number of these tissues are involved in resistance to pathogen pressure (e.g. blood, esophageal mucosa, and the skin), and there is prior evidence that immune response is an important selective pressure [62]. We note eQTLs that affect sun-exposed skin are more likely to be adaptive outliers than eQTLs that affect skin that has not been exposed to the sun, and there is evidence that sun-exposure has contributed to adaptation [63]. Many tissues that are not exposed to external environments have a relative lack of adaptive outliers (Figure 4). We also find that eQTLs that modify gene expression in the cerebellum are more likely to be adaptive than eQTLs that modify gene expression in other brain tissues. This pattern is consistent with morphological evidence that cerebellum volumes have increased more than neocortex volumes during the evolution of humans and other great apes [64].

Focusing on sex-specific tissues, we note that the effective strength of selection is halved if an eQTL only affects expression in one sex. For example, testis-specific eQTLs contribute to male, but not female, fitness. This is offset by the potential of these tissues to be targets of sexual selection. We find that eQTLs that affect male-specific tissues (testis and prostate) are more likely to be adaptive than eQTLs that affect female-specific tissues (uterus, ovary, and vagina). What might contribute to accelerated evolution of male eQTLs? One contributing factor is that eQTLs that affect female tissues tend to be more pleiotropic, i.e. they affect a larger number of additional tissues. There is also genetic evidence that spermatogenesis has been a key target of positive selection in human evolution [65]. An additional contributing factor is that variability in reproductive success is greater for males than females (Bateman’s principle) [66]. This suggests that that the accelerated evolution of testis eQTLs may be driven by increased sexual selection acting on males.

Our study focused on evolution that has occurred in the last 100,000 years. Different eQTLs and tissues may have been locally adaptive over different timescales. That said, the patterns seen in our study mirror what is seen other primates: adaptive genetic changes are more likely to affect gene expression in testes than the brain [67]. There is also evidence that some regions of the genome are recurrent targets of adaptation (20q13, for example). Consistent with theoretical predictions [68], we find that most adaptive alleles do not have large effect sizes and that pleiotropy appears to inhibit adaptation. Here, we focused on positive selection, as opposed to other forms of selection. We note that gene expression is a trait that is often under stabilizing selection [69]. In addition, eQTLs can be subject to negative selection. Negative selection acting directly on eQTLs can result in lower values of F_ST_. However, negative selection at linked sites reduces effective population sizes – causing F_ST_ statistics to be slightly elevated for genomic regions that experience background selection [70]. Regardless, GTEx and RegulomeDB eQTLs with very large F_ST_ statistics and PBS scores are best explained by positive selection.

The findings described here have implications for health and disease. Many GWAS risk alleles influence gene expression [59, 71, 72]. Furthermore, a large number of adaptive eQTLs are closely linked to loci that contribute to complex traits and disease risks in human populations. Because PBS outliers have highly divergent allele frequencies between populations, the combination of positive selection on gene expression and genetic hitchhiking can contribute to health inequities. In addition, hereditary disease risks have evolved over time [24], with a major contributing factor to this being changes in gene expression. One implication is that diseases affecting tissues that are enriched for PBS outliers are more likely have disease risks that have changed greatly over time.

One caveat of our study is that the set of presently-known eQTLs is only the tip of the iceberg. Because the statistical power to detect eQTLs depends on effect size [73], many eQTLs with small effects sizes have yet to be detected. We also note that human eQTLs have largely been ascertained in individuals that have European ancestry. Despite this bias, similar patterns are observed for eQTLs that have signatures of positive selection in African, European, or East-Asian individuals (Figures S3-S5. Furthermore, we note that the GTEx project was able detect eQTLs like rs2814778, where the adaptive allele is rare outside of Africa. Many putatively adaptive alleles are actually eQTLs that have not yet been identified as such (how else to explain noncoding variants that have such divergent allele frequencies across populations?). For example, the European PBS peak at 15q21.1 contains SNPs that affect pigmentation, but these SNPs were not classified as eQTLs because they did meet our stringent p-value filter of 10^-9^.

An additional consideration is that rare alleles are underrepresented in the set of known eQTLs (Figure S9). This occurs because the statistical power to detect an eQTL is maximized when SNPs have large minor allele frequencies [74]. SNPs with large minor allele frequencies are also more likely to be high F_ST_ SNPs. Because of this, many presently known eQTLs have high F_ST_ statistics (Figure S10). Differences in genotyping technologies cause V7 GTEx eQTLs to have F_ST_ distributions that are left-shifted compared to V6 GTEx eQTLs [75, 76]. With the exception of the lowest minor allele frequency bin, frequency-matched eQTLs have F_ST_ statistics that are similar to random SNPs from the 1000 Genomes Project (Figure S9). As sample sizes increase in the future, additional rare eQTLs will be able to be discovered. However, these rare eQTLs are unlikely to have high values of F_ST_ [77]. Finally, we note that highly stringent PBS cutoffs were used to identify adaptive eQTLs. For example: allele frequencies of 60%, 10%, and 10% give PBS scores that are below the thresholds shown in Figure 1.

Going forward, future studies will lead to the discovery of eQTLs that affect additional tissues, examine whether decreasing or increasing expression is more likely to be adaptive, and identify the extent that eQTLs are found in adaptively introgressed haplotypes. In conclusion, we find that many eQTLs have been positively selected, and these adaptive eQTLs reveal important details about the recent evolution of our species.

## Material and methods

### Population genomic data and eQTL datasets

Allele frequencies at 80,701,406 autosomal SNPs were obtained from Phase 3 of the 1000 Genomes Project [78]. Continental super-populations from the 1000 Genomes Project were used: Africa (ACB, ASW, ESN, GWD, LWK, MSL, and YRI), Europe (CEU, FIN, IBS, GBR, and TSI), and East Asia (CDX, CHB, CHS, KHV and JPT). Sample sizes varied for each continent population: 661 individuals of African descent 503 individuals of European descent, and 504 individuals of East Asian descent. Biallelic SNPs from Phase 3 of the 1000 Genomes Project (ascertained via whole genome sequencing) were merged with rs # identifiers from the Illumina Omni 2.5M array, RegulomeDB, and Genotype-Tissue Expression (GTEx) Project. RegulomeDB scores of 1a, 1b, 1c, 1d, 1e, or 1f indicate that a SNP is a RegulomeDB eQTL [17]. For GTEx eQTLs, we required sample sizes of at least 70 individuals per tissue, yielding 44 tissues for V6 and 48 tissues for V7. To correct for multiple statistical tests, V7 GTEx eQTLs were required to have a p-value ≤ 10^-9^ for at least one tissue. Allele frequency and eQTL data were merged using the *dplyr* package in R [79]. Genomic positions described here are from the GRCh37/hg19 assembly. Most analyses in this paper focused on V7 GTEx eQTLs. SNPs were also binned into 10% minor allele frequency (MAF) bins. European MAF bins were used because the majority of eQTLs have been ascertained in individuals that have European ancestry.

### Genetic distances and scans of selection

Weir and Cockerham’s F_st_ was calculated for each pairwise combination of populations (AFR-EUR, AFR-EAS, and EUR-EAS) [77, 80]. This method of calculating genetic distances corrects for small sample sizes. Five types of SNPs were analyzed: SNPs from the 1000 Genomes Project (ascertained via whole genome sequencing), SNPs on the Illumina Omni 2.5M array, V6 and V7 GTEx eQTLs, and RegulomeDB eQTLs. Empirical cumulative distribution functions and mean values of F_st_ and were found for each type of SNP and population pair. To identify adaptive SNPs, population branch statistics (PBS) [36, 37] were then calculated for V6 and V7 GTEx eQTLs using the following equations:

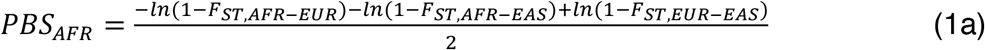

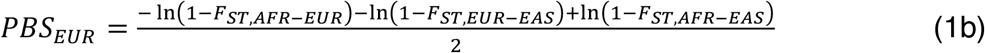

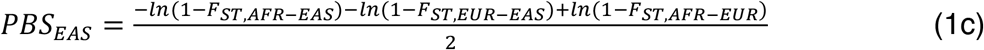

Undefined and negative values of Weir and Cockerham’s F_st_ were treated as zero for PBS calculations. Genome-wide distributions of PBS scores were calculated for each branch (Africa, Europe, and East Asia). In total, this yielded 1,826,392 PBS scores for V6 GTEx eQTLs and 1,154,731 PBS scores for V7 GTEx eQTLs. Negative PBS scores were treated as zero. Adaptive eQTL outliers were required to have PBS scores above the 99th percentile for a particular branch. To correct for the effects of linkage disequilibrium, we selected the eQTL with the top PBS score in each 100kb genomic window. This yielded a total of 614 (African), 561 (European), and 524 (East Asian) LD-pruned adaptive eQTL outliers. Due to greater amounts of linkage disequilibrium [78], high PBS eQTLs tend to cluster more in non-African genomes. This causes the African branch to have a larger number of LD-pruned adaptive eQTLs. An LD-pruned cutoff of the top 1% yields enough PBS outliers per tissue to examine the tissue-specificity of adaptation. As an additional control, we repeated our analyses of adaptive outliers using a cutoff of the top 0.1% PBS scores (LD-pruned). However, we note that using a top 0.1% PBS score cutoff prevents accurate inference of tissue-specific enrichment ratios (this is because 19 tissues have less than 10 LD-pruned adaptive outliers that meet this stringent threshold).

### Pleiotropy and tissue breadth

Highly pleiotropic eQTLs modify expression a large number of tissues. We generated tissue breadth scores for each eQTL by finding the number of tissues with p-values ≤ 10^-9^. This yielded tissue breadth scores between 1 and 48 for V7 GTEx eQTLs. To identify whether adaptive eQTLs affect a different number of tissues than non-adaptive eQTLs, empirical cumulative distribution functions and mean values of tissue breadth scores were found for non-adaptive eQTLs and adaptive PBS outliers.

### eQTL effect sizes

We tested whether eQTLs with large allele frequency differences between populations also yield large differences in gene expression. Here, we focused on LD-pruned eQTLs (i.e. eQTLs with the highest PBS score per 100kb window) that only affect a single tissue. Effect sizes for each eQTL and tissue combination were quantified by taking the absolute value of normalized effect size (NES) under a fixed effect model, i.e. |β_FE_| For each LD-pruned GTEx eQTL, PBS scores were plotted against |β_FE_|. The “ggscatter” function in the *ggpubr* R package was used for local regression (loess) fitting and to assess the extent to which PBS scores and |β_FE_| values are correlated. Note that loess fitting allows non-linear relationships to be identified. This analysis was repeated for three different continental populations (African, European, and East Asian) and all 48 tissues in the V7 GTEx dataset. Wilcoxon rank sum tests were used to determine whether single tissue eQTLs with PBS scores in the top 1% have effect sizes that differ from eQTLs with PBS scores in the bottom 99%.

### Tissue-specific adaptation

Here, PBS outliers are LD-pruned eQTLs that have PBS scores in the top 1% of all GTEx eQTLs. The proportion of eQTLs that are LD-pruned PBS outliers was found by taking the geometric mean the number of outliers divided by the number of eQTLS for each tissue:

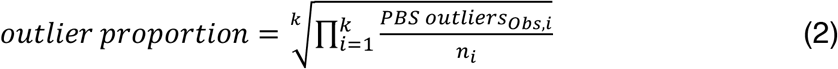

where *k* is the total number of tissues and *n* is the total number of eQTLs for tissue *i*. We then found the expected number of adaptive PBS outliers for each tissue:

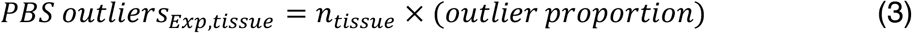

The number of adaptive PBS outliers was compared to the total number of eQTLs for each tissue. This allowed us to identify the primary targets of adaptive evolution. Enrichment ratio statistics were calculated for each tissue by comparing the observed number of adaptive PBS outliers to the expected number of adaptive PBS outliers:

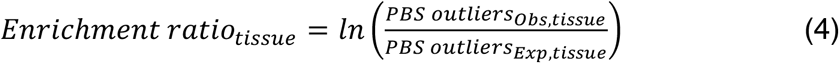

The geometric mean used in Equation 2 ensures that the sum of all tissue-specific enrichment ratios generated using Equation 4 is zero. Positive enrichment ratios indicate tissues with a relative excess of adaptive eQTLs, and negative enrichment ratios indicate that tissues with a relative lack of adaptive eQTLs. Enrichment ratio statistics were calculated for 48 V7 GTEx tissues and 44 V6 GTEx tissues. 95% confidence intervals for enrichment ratios were found by using the Agresti-Coull approach to find lower and upper bounds for tissue-specific outlier proportions [81]. We also adjusted for sample size as covariate via the following equation:

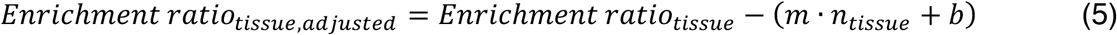

where *n_tissue_* is the number of individuals sampled per tissue, and *m* and *b* are the slope and intercept of the regression line in Figure S7 (i.e. *m* = 0.0012 and *b* = -0.2517). Adjusted enrichment ratios measure how much a given tissue is above or below the regression line in Figure S7, i.e. they are the residuals.

### Overlap with GWAS loci

We assessed overlap between adaptive eQTLs and GWAS loci by downloading data from the NHGRI-EBI GWAS Catalog [82, 83]. For this analysis, LD-pruned sets of V7 GTEx eQTLs with PBS scores in the top 1% were considered to be adaptive. Distances between adaptive eQTLs and GWAS loci were found using the “closest” function in BEDTools [84].

## Acknowledgements

We thank Greg Gibson, Michelle Kim, Urko Martinez-Marigorta, Corinne Simonti, and Biao Zeng for helpful comments and suggestions. This work was supported by startup funds from Georgia Institute of Technology. Melanie Quiver was also supported by an NIH training grant (T32GM105490).

## Supplemental Material

**Figure S1.**
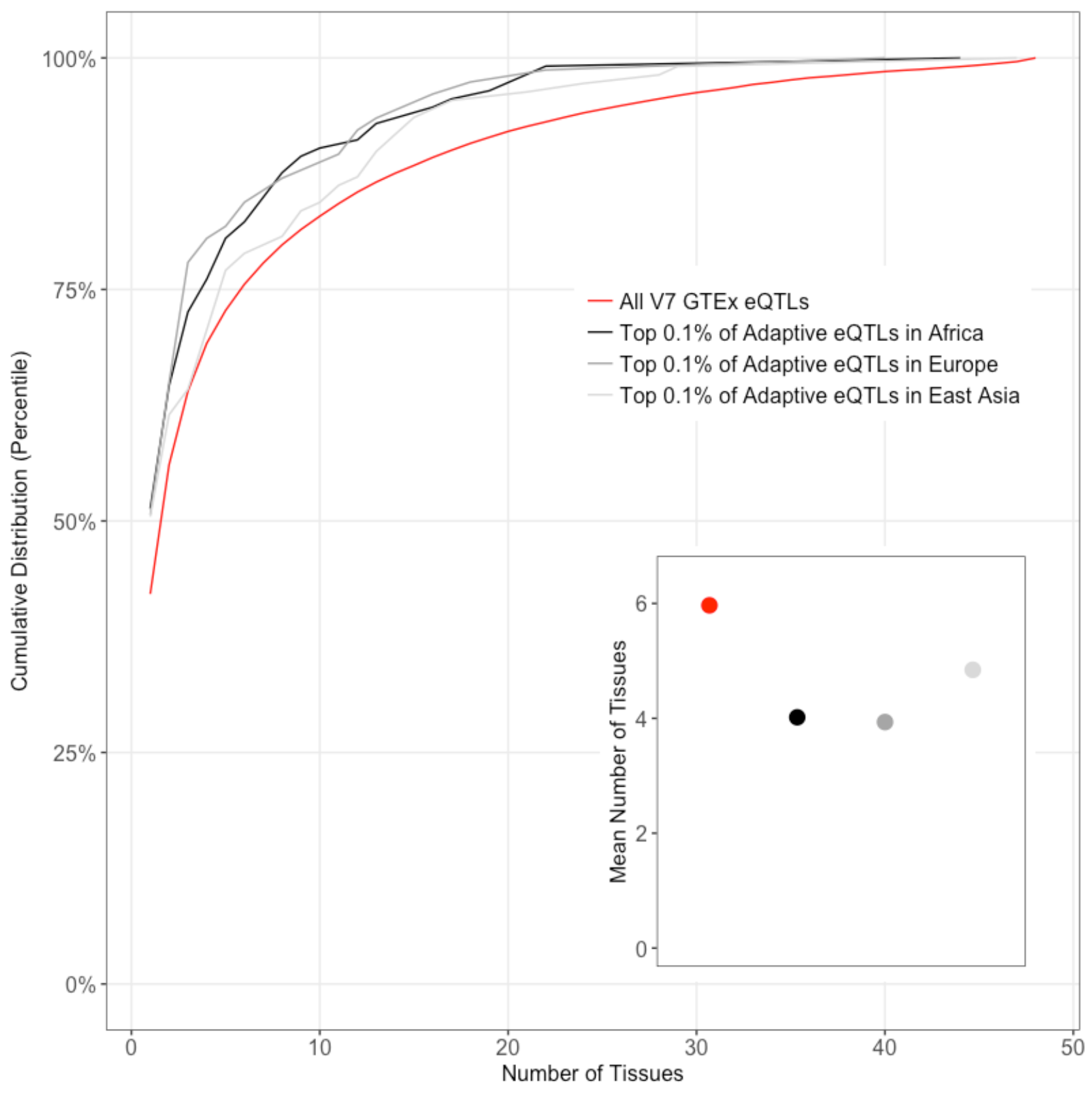
Pleiotropy results are robust to the use of a more stringent adaptive outlier threshold. Here, adaptive outliers are LD-pruned eQTLs that have PBS scores in the top 0.1% of all GTEx eQTLs. Cumulative distributions and mean number of tissues are shown for adaptive and non-adaptive eQTLs. In general, adaptive eQTLs modify gene expression in fewer tissues than non-adaptive eQTLs. For each population, differences in the number of tissues affected by adaptive outliers and the overall set of GTEx eQTLs are statistically significant (p-value < 2.2 10-16 for all comparisons, Wilcoxon rank sum tests).

**Figure S2.**
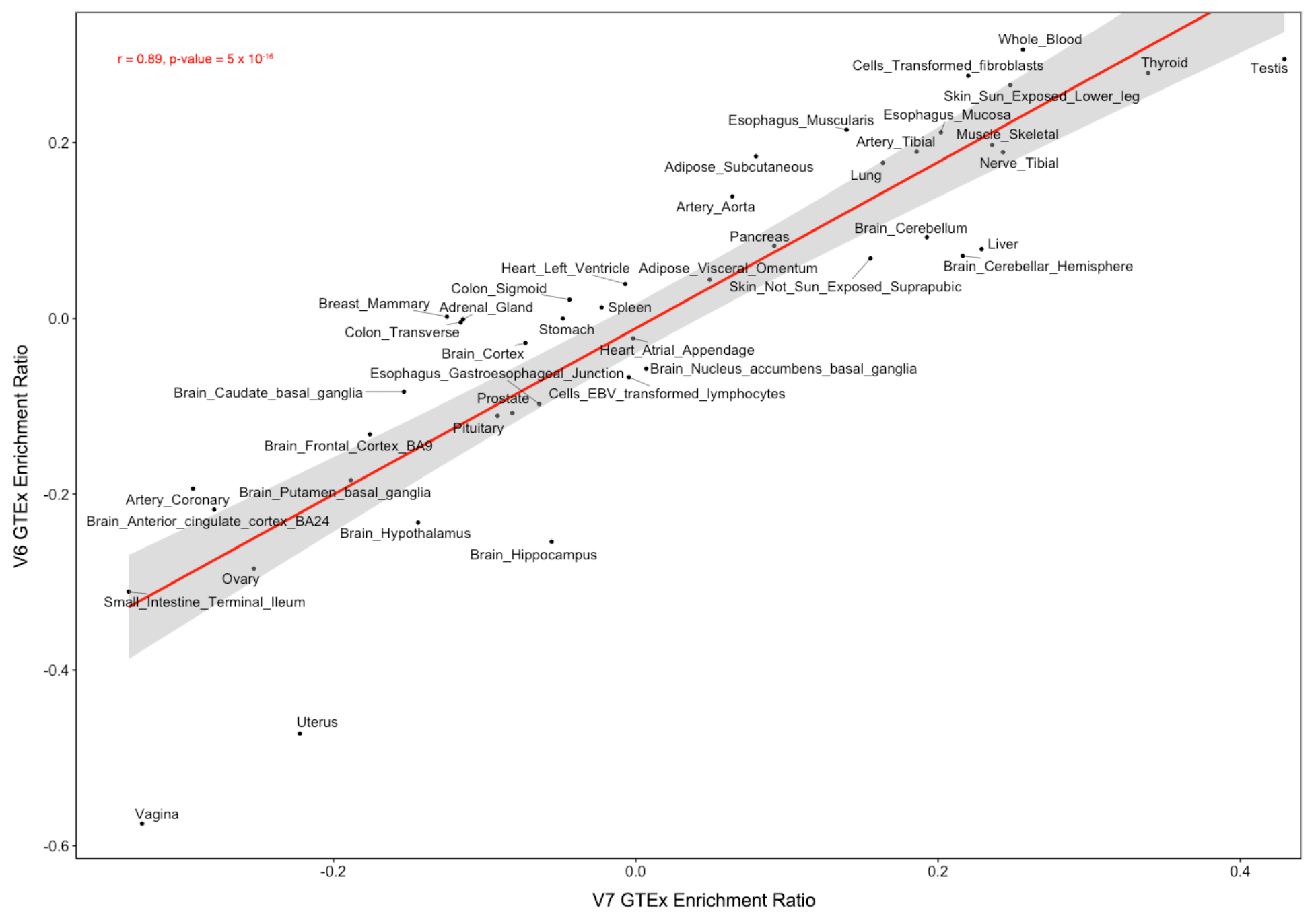
Enrichment ratios are similar for different versions of GTEx. 44 tissues have data for V6 and V7 of GTEx. Each data point is a single tissue. Positive enrichment ratios indicate a relative excess of adaptive eQTLs and negative enrichment ratios indicate a relative lack of adaptive eQTLs.

**Figure S3.**
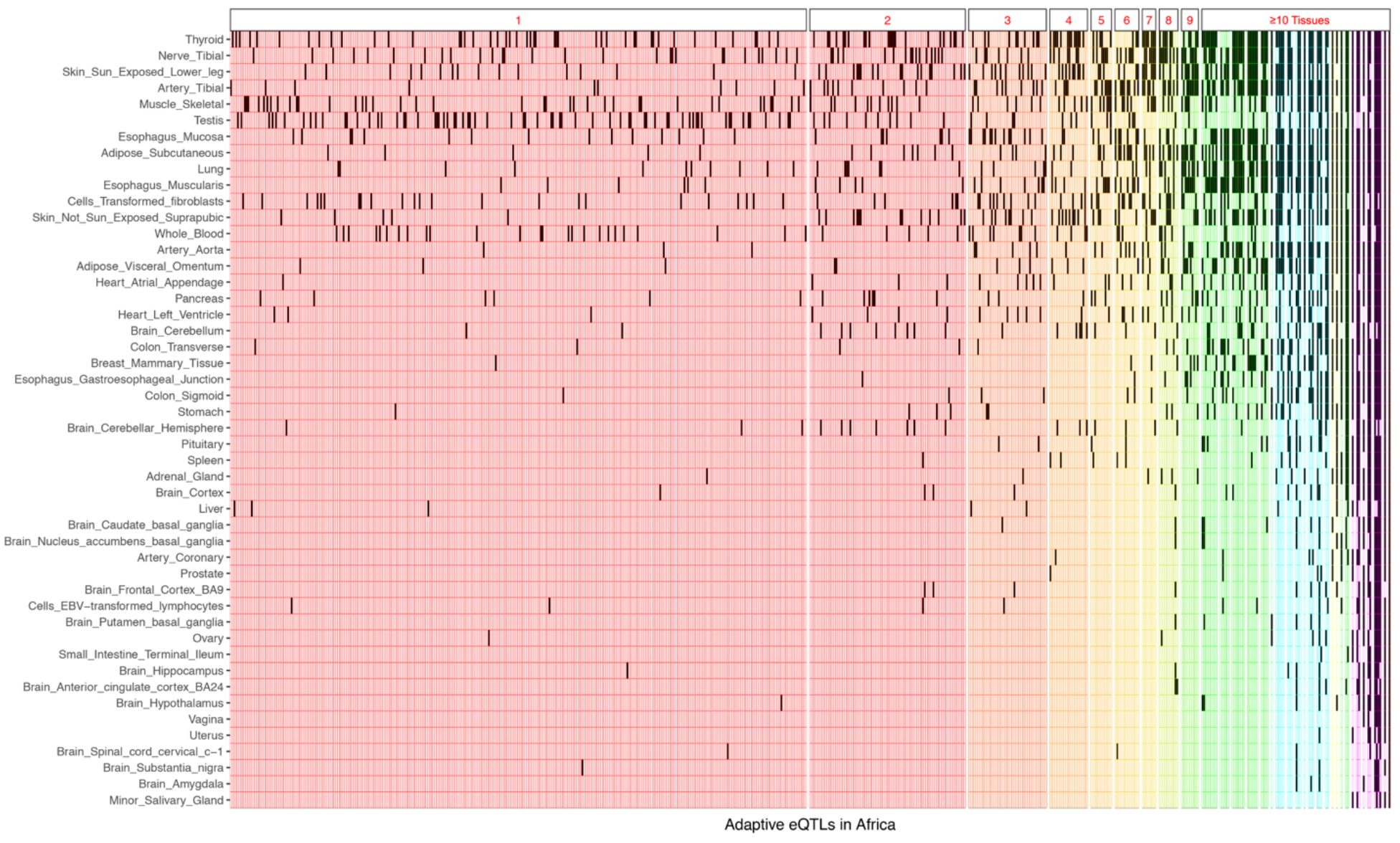
Tissue-specificity of African PBS outliers. A grid of 48 rows and 614 columns is shown, where each row corresponds to a different tissue and each column corresponds to a different adaptive eQTL for the African branch. Filled cells reveal which tissues are affected by each adaptive eQTL. Background colors indicate the number of tissues affected by each adaptive eQTL (e.g. red indicates eQTLs that modify expression in a single tissue). Tissues are rank-ordered by total number of adaptive eQTLs.

**Figure S4.**
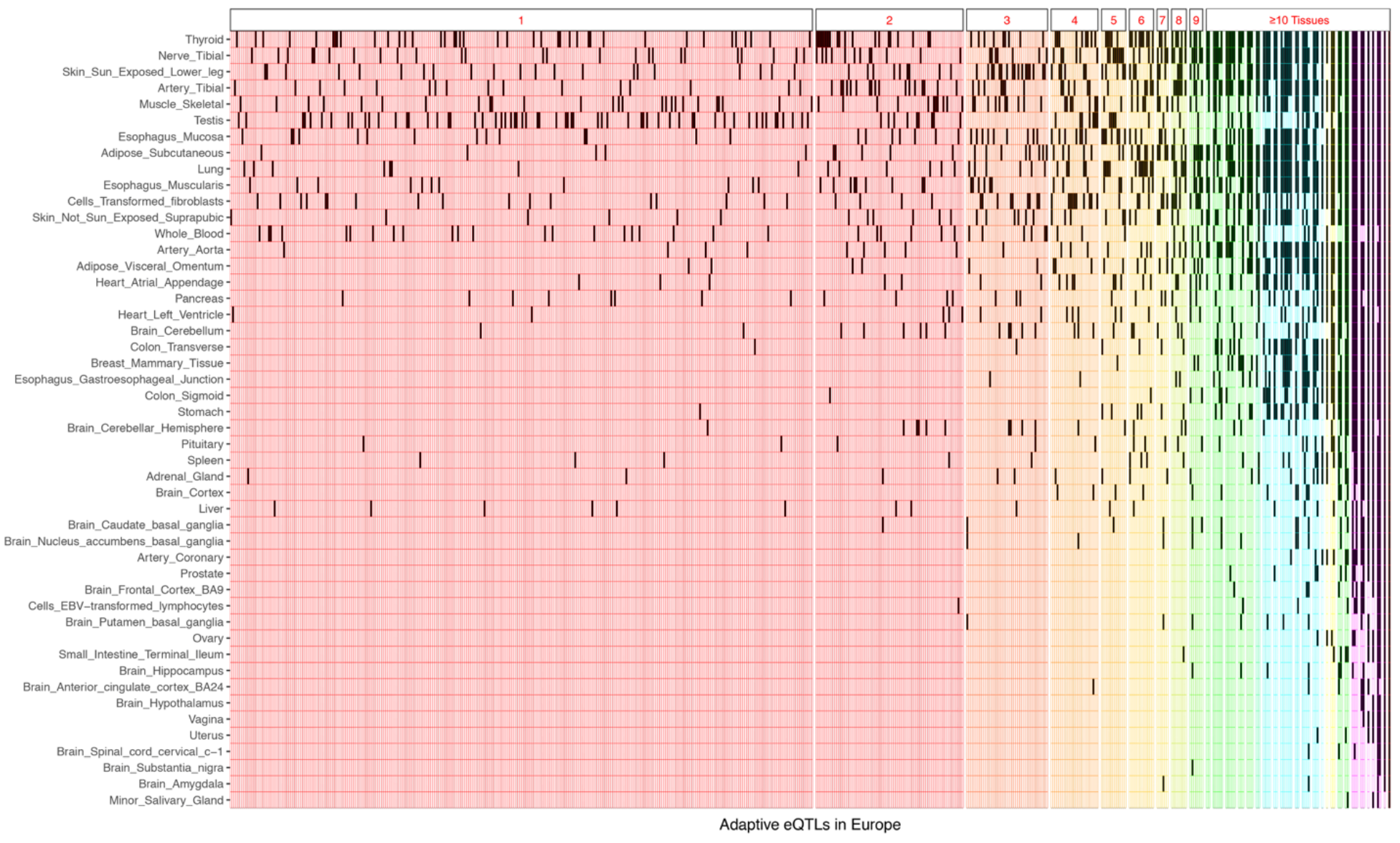
Tissue-specificity of European PBS outliers. A grid of 48 rows and 561 columns is shown, where each row corresponds to a different tissue and each column corresponds to a different adaptive eQTL for the European branch. Filled cells reveal which tissues are affected by each adaptive eQTL. Background colors indicate the number of tissues affected by each adaptive eQTL (e.g. red indicates eQTLs that modify expression in a single tissue). Tissues are rank-ordered by total number of adaptive eQTLs.

**Figure S5.**
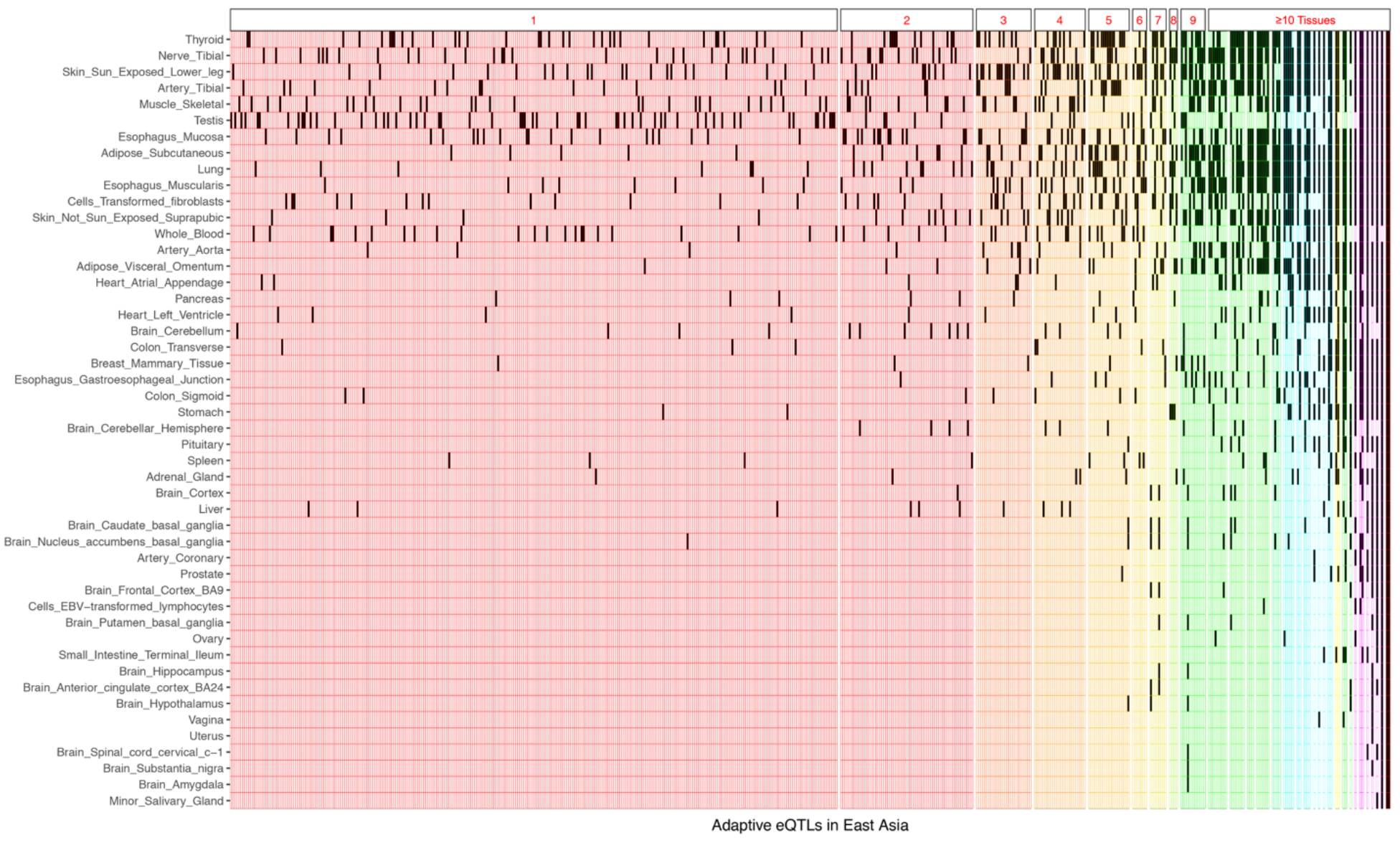
Tissue-specificity of East Asian PBS outliers. A grid of 48 rows and 524 columns is shown, where each row corresponds to a different tissue and each column corresponds to a different adaptive eQTL for the East Asian branch. Filled cells reveal which tissues are affected by each adaptive eQTL. Background colors indicate the number of tissues affected by each adaptive eQTL (e.g. red indicates eQTLs that modify expression in a single tissue). Tissues are rank-ordered by total number of adaptive eQTLs.

**Figure S6.**
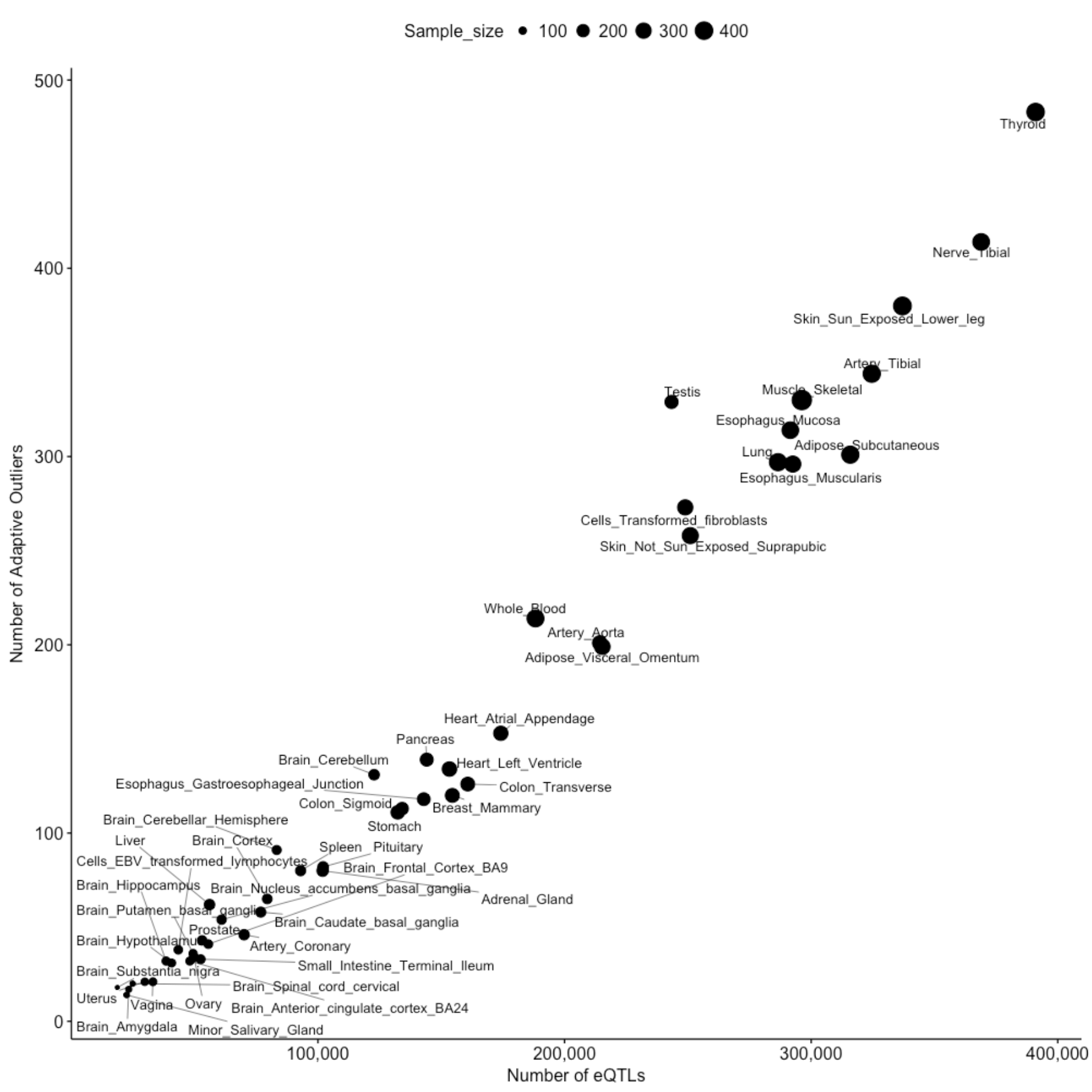
Numbers of known eQTLs vary by tissue. Each point represented a different tissue, with the size of each point indicating sample size (numbers of individuals). Here, adaptive outliers are LD-pruned eQTLs that have PBS scores in the top 1% of all V7 GTEx eQTLs. Larger sample sizes result in a greater number of eQTLs being detected.

**Figure S7.**
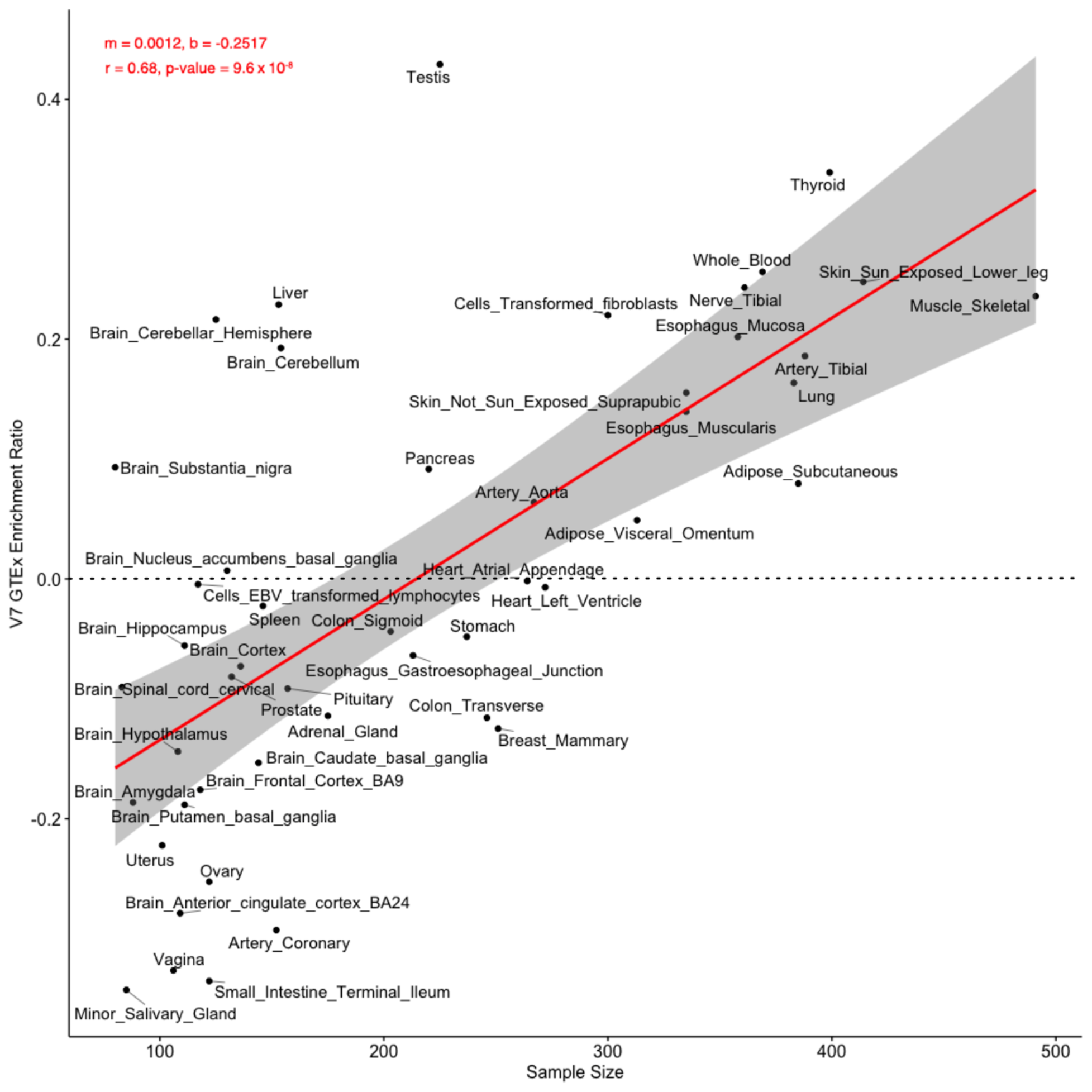
Enrichment ratios plotted against sample sizes per tissue. Equation for the regression line: Enrichment Ratio = 0.0012 x (Sample Size) – 0.2517. Tissues with positive adjusted enrichment ratios are above the solid red line, and tissues with negative adjusted enrichment ratios are below the solid red line.

**Figure S8.**
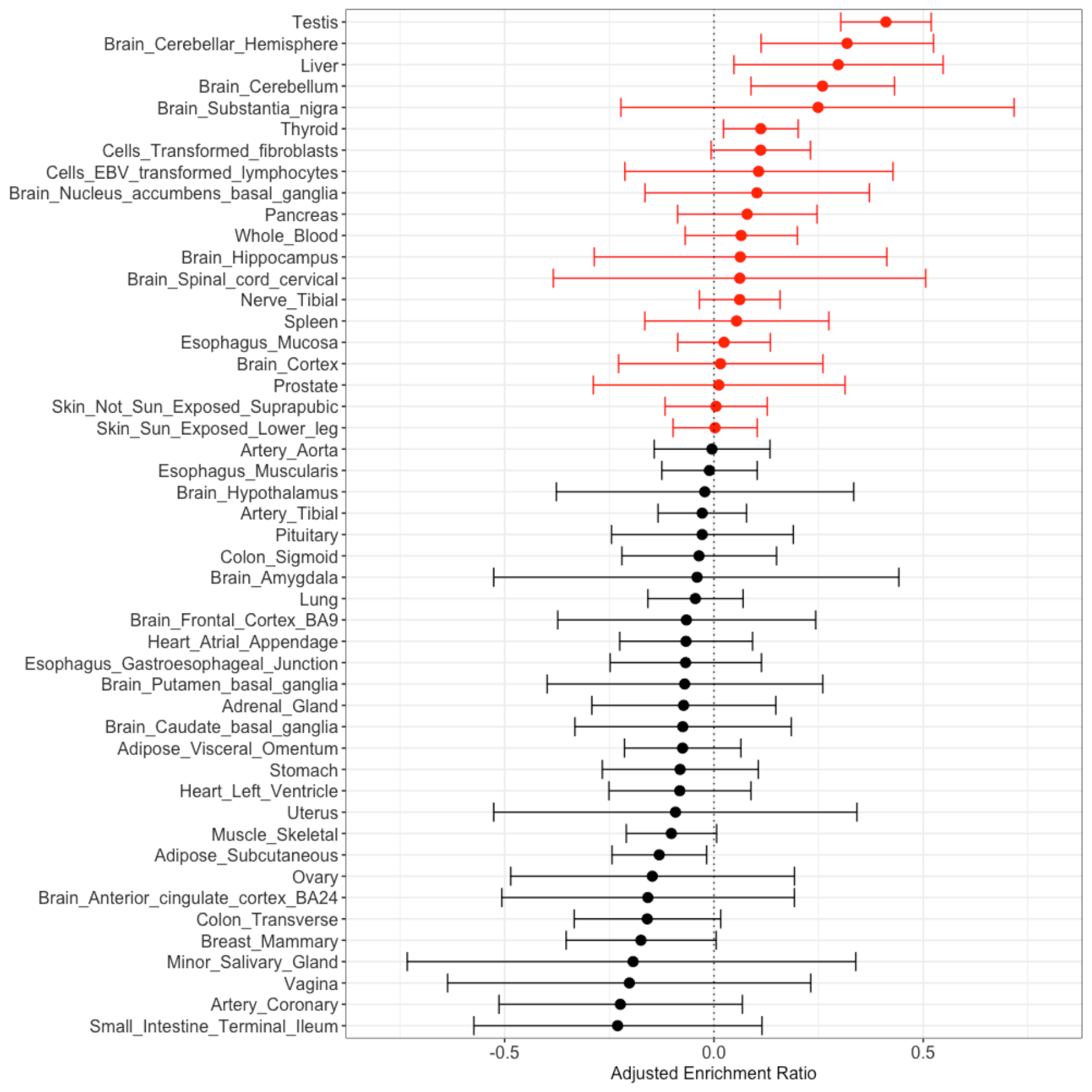
Tissue enrichment results after correcting for sample size differences using a linear model. Here, adaptive outliers are LD-pruned eQTLs that have PBS scores in the top 1% of all GTEx eQTLs. 95% confidence intervals for each adjusted enrichment ratio are shown. Adjusted enrichment ratios are the residuals from Figure S7.

**Figure S9.**
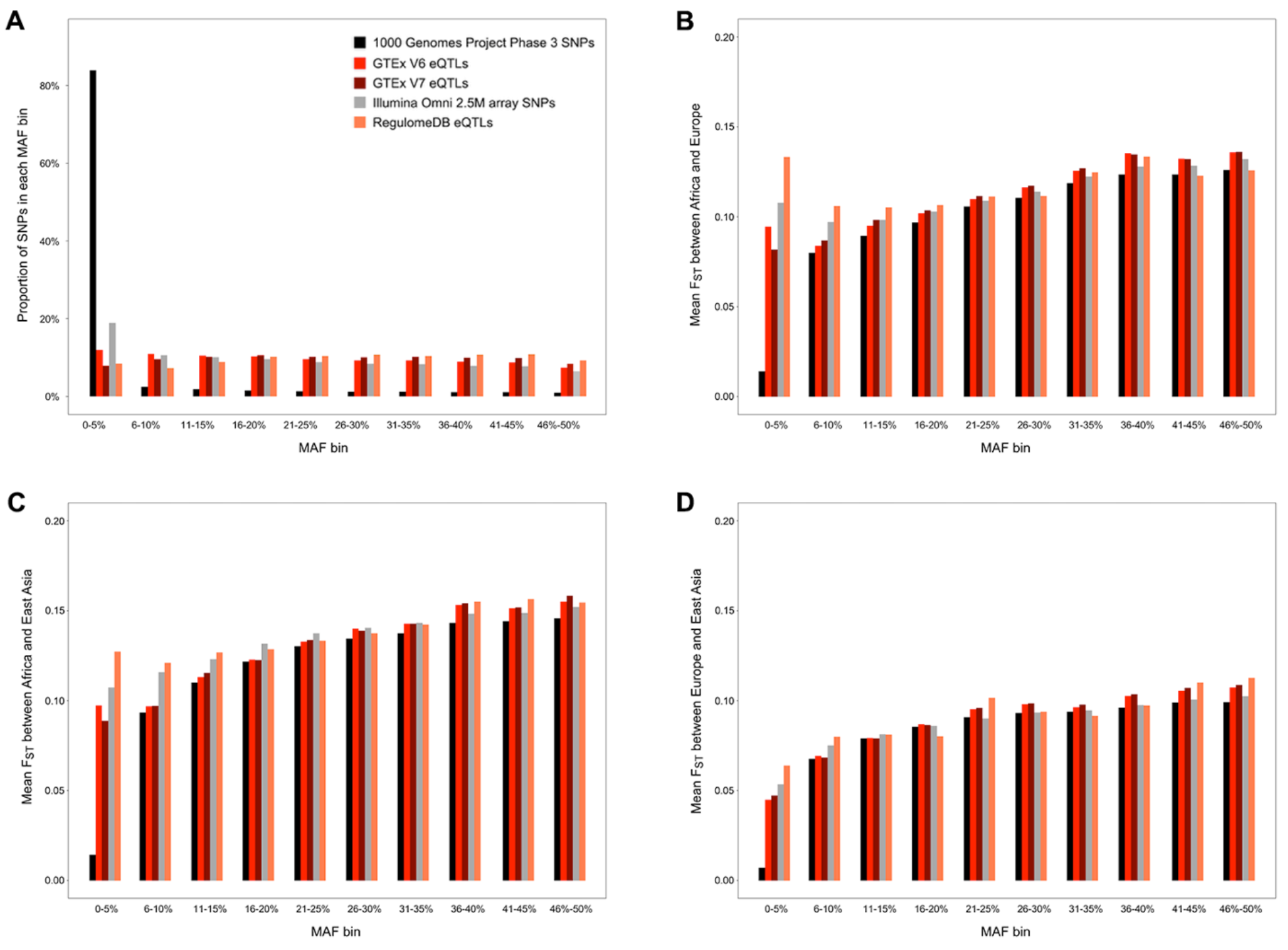
Genetic distances between populations vary by minor allele frequency (MAF). Panel (A) shows the proportion of each type of SNP that is found in each European MAF bin. Conditioning on MAF bin, mean F_ST_ values between different pairs of populations are shown for different types of SNPs in panels (B) African and European populations, (C) African and East Asian populations, and (D) European and East Asian populations.

**Figure S10.**
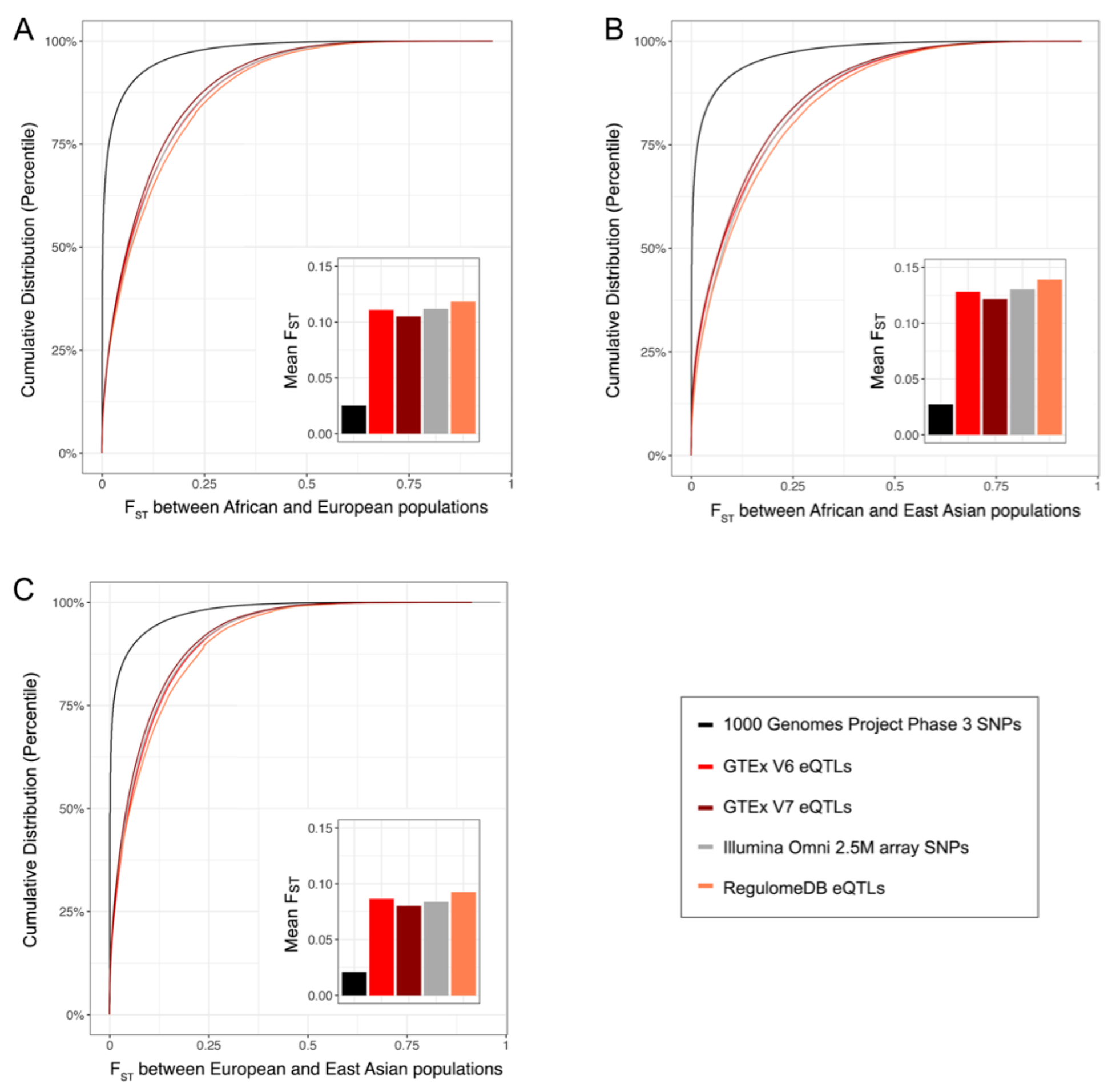
Presently known eQTLs tend to have higher values of F_ST_ statistics than random SNPs from the 1000 Genomes Project. Genetic distances between different pairs of populations were calculated using Weir and Cockerham’s F_ST_: (A) African and European populations, (B) African and East Asian populations, and (C) European and East Asian populations. Cumulative distributions and mean values of F_ST_ are shown for each type of SNP.

